# Assessment of cognitive reserve using near infrared spectroscopy

**DOI:** 10.1101/276865

**Authors:** Andrei V. Medvedev

## Abstract

Cognitive reserve (CR) is the ability to preserve cognitive functions in the presence of brain pathology. One commonly used proxy measure of CR is IQ. In the context of Alzheimer’s disease (AD), patients with higher CR show better cognitive performance relative to brain damage therefore higher CR reduces the risk of dementia. There is a strong need to develop a reliable biomarker of CR given the growing interest in understanding protective brain mechanisms in AD. Recent fMRI studies indicate that frontoparietal network may play an important role in the maintenance of cognitive reserve. The goal of this study was to measure functional connectivity (FC) of the prefrontal cortex using near infrared spectroscopy (NIRS) and to study the relationship of prefrontal FC with cognitive abilities and motoric skills.

We analyzed resting state optical data recorded from prefrontal cortex in 13 healthy individuals who were also assessed by Wechsler Abbreviated Scale of Intelligence (WASI) test and the Purdue Pegboard test (PPT). For each participant, activity of each prefrontal channel was correlated with all other channels and positive correlation coefficients were Fisher-transformed and averaged over all PFC channels giving the Global Functional Connectivity (GFC) of PFC. The resulting GFC number for each individual was then correlated with the corresponding IQ (WASI full score) and the PPT scores.

Prefrontal connectivity was found to positively correlate with IQ while showing the lack of or negative correlation with the Purdue Pegboard subtests. These results demonstrate that the cost-effective and noninvasive NIRS technology can be used to evaluate prefrontal functional connectivity thus providing a neurophysiological measure of cognitive reserve.

E-mail address for correspondence:

## 1. Introduction

Cognitive processes and their changes in healthy and pathological aging have been investigated in detail over the past few decades which led to the development of new theoretical models of aging. Existing data show a complex picture of age-related changes in cognitive functions, which can include both the relative stability and deterioration. Thus, many perceptual and cognitive processes demonstrate a general slowing e.g., leading to difficulties in finding the right word during speech production. Age- and pathology-related microstructural changes in gray and white matter may underlie the observed reduction in processing speed, working memory capacity and hearing loss which reduce cognitive abilities (reviewed in Burke and Mackay, 1997). Nevertheless, despite those deficits, older people may show a relative stability of certain cognitive functions despite the observed microstructural pathology. This suggests the existence of both dysfunction and possibly compensation in functional networks (for review, see Stern, 2009) and naturally raises the question of substitute or compensatory mechanisms by which this stability, commonly referred to as cognitive reserve (CR), is achieved.

The concept of CR was introduced by Stern to explain the absence of a direct relationship between the degree of brain pathology or damage and the clinical manifestation of that damage (Stern, 2009). For example, in several studies it has been demonstrated that up to 25% of senior people have unimpaired cognitive functioning revealed by neuropsychological testing prior to death while meeting full pathologic criteria for AD (for review, see Stern, 2009). This suggests that this degree of pathology does not necessarily lead to clinical dementia. The concept of cognitive reserve posits that individual differences in functioning of neural networks allow some people to withstand the pathological process and maintain a relatively high level of cognitive functions. The CR concept is supported by numerous epidemiological studies. For example, Valenzuela and Sachdev in their meta-analysis reviewed 22 papers reporting cohort studies of the effects of education, occupation, premorbid IQ and mental activities in incident dementia (Valenzuela and Sachdev, 2006). Ten out of 15 studies revealed a significant protective effect of education; 2 out of 2 showed a protective effect of premorbid IQ; 9 out of 12 demonstrated a protective effect of occupational attainment and 6 out of 6 showed a protective effect of engaging in leisure activities (reviewed in Stern, 2009). The authors concluded that higher CR (as measured by IQ and years of education) was significantly correlated with a lower risk for incident dementia.

Different neural mechanisms may underlie cognitive reserve and Stern (2009) suggested two major components, neural reserve reflecting just individual differences in neural networks and neural compensation which requires alterations in cognitive processing in order to cope with brain pathology. Thus, a more efficient use of brain networks and/or recruitment of alternate brain networks may lead to the development of CR. Numerous studies indicate that environmental enrichment and specific life experiences such as educational activities and occupation play an important role in developing CR and reducing risk of dementia (Stern, 2009).

*Measures of cognitive reserve*. To assess CR, various measures of socioeconomic status are used such as occupational, educational (including degree of literacy) and leisure activity. Many studies suggest that IQ is a powerful measure of reserve because it reflects innate intelligence (Albert and Teresi, 1999).

Evidence is emerging that functional architecture of the brain is relatively stable in a variety of cognitive tasks and the resting state. Recent brain imaging studies have demonstrated the existence of several networks within the brain which can be distinguished by correlated activity of their nodes over time within each network and uncorrelated or anti-correlated activity between networks. Those networks have been suggested to represent an essential feature of the functional architecture of human brain (Fox et al., 2005). Thus, their analysis is important for our understanding of normal and impaired cognitive functions. This network organization seems to exist independently of the functional state of the brain because the same patterns of correlated and anti-correlated activity are observed during cognitive task performance as well as in the absence of any task or stimulus i.e., in the resting state (Fox et al., 2005; Fox and Raichle, 2007). Slow (<0.1 Hz) fluctuations of brain activity correlated within the network can be recorded by measuring BOLD fMRI signal (for review, see Fox and Raichle, 2007). Similar signals can be detected in blood hemoglobin levels when measured by functional near-infrared spectroscopy (NIRS) (Strangman et al., 2002; Medvedev et al., 2011; Medvedev, 2013). Networks of brain regions that are activated during cognitive tasks (e.g., working memory) have been labeled as “task-positive” (TPNs). Other networks, by contrast, showing deactivation during cognitive tasks but activation in the resting state have been labeled as “task-negative” (TNNs or the “default mode network”, DMN). The TPNs and TNNs show either no correlation or even anti-correlative relationship to each other that is observed during both task performance and the resting state (Fox et al., 2005).

The strength of functional interactions within and between networks (network connectivity) has been suggested to be one of the key determinants of cognitive abilities in general and more specifically, cognitive control which is the ability to effectively control thoughts and behavior in everyday life. Cognitive control can be measured by working memory capacity and general fluid intelligence (Cole et al., 2012). The general feature of cognitive control is its limited capacity which predicts important life relevant outcomes such as academic and professional success (Cole et al., 2012). The fundamental property of cognitive control is its extraordinary ability to adaptively organize a wide variety of tasks thus providing a central mechanism allowing an individual to switch between different tasks in a highly flexible manner as well as to learn new tasks. There is an emerging consensus that there is a core set of brain structures which is centrally involved in cognitive control – a *cognitive control network*. This functional network consists of the lateral prefrontal cortex (LPFC) and the posterior parietal cortex (PPC) making up the so-called frontoparietal network (FPN), and it has been variously termed the cognitive control network or system (Cole et al., 2012), the superordinate cognitive control network (Niendam et al., 2012), the multiple-demand system (Duncan, 2010) and the task-positive network (Fox et al., 2005). The frontoparietal network has received a lot of attention in recent studies (Cole et al., 2012; Cole et al., 2013; Franzmeier et al., 2017).

The goal of this study was to demonstrate that near-infrared spectroscopy can be used to measure functional connectivity of the lateral prefrontal cortex and to explore whether this connectivity correlates with cognitive and behavioral performance specifically, the IQ, a proxy measure of cognitive reserve, as well as the Purdue Pegboard Test (PPT) in healthy individuals.

## 2. Methods

In this study, we used the resting state data from the same cohort of adult subjects which were previously analyzed in the context of hemispheric asymmetry of functional connectivity (Medvedev, 2013). The details of the methods can be found there. Briefly, 13 right-handed subjects reported as being in good health, having normal (or corrected to normal) vision and without medications and signed a consent form approved by the Georgetown University Institutional Review Board. Before experiments, they undertook a battery of cognitive and behavioral tests which included measures of IQ (Wechsler Abbreviated Scale of Intelligence), handedness and Purdue Pegboard Test which is commonly used for finger and hand dexterity and bimanual coordination. Subjects were scanned during 4-8 min rest before engagement in a cognitive task. Task-related data have been published elsewhere (Medvedev et al., 2011) and for the current study, the resting state data were analyzed. Optical signals were recorded using a continuous-wave NIRS instrument CW5 (TechEn, Milford, MA) with two 14 × 8-cm probes, each accommodating 11 optodes with three dual-wavelength (690 and 830 nm) laser sources and eight detectors for each hemisphere. Optical sources and detectors were placed on the scalp using the 10-20 EEG electrode placement system with locations F3/4-F7/8-C3/4 used as reference points. As a result, optical probes were covering inferior frontal gyrus (IFG) and the middle frontal gyrus (MFG), the essential components of the frontoparietal cognitive control network.

For each participant, we calculated the Global Functional Connectivity (GFC) of the lateral prefrontal cortex following the procedure described in (Cole et al., 2012) adapted for our optical data. Briefly, GFC was determined first for each optical channel covering the left and right lateral PFC (following optode localization procedures as in Fishburn et al., 2014). Brain activity was analyzed separately for oxygenated hemoglobin (HbO), deoxygenated (reduced) hemoglobin (HbR) and total hemoglobin (HbT = HbO + HbR). Activity of each prefrontal channel was correlated with all other channels and all positive correlation coefficients were z-transformed (by Fisher’s transform) and averaged over all PFC channels (as per Cole et al., 2012 and Franzmeier et al., 2017). For each individual, the resulting GFC numbers (separately for HbO, HbR and HbT) were then correlated with his/her IQ (WASI full score) and two scores of the two PPT subtests namely, the score for the performance with both hands (PP Both Hands) and the score for the assembly subtest (PP Assembly).

## 3. Results

First, we divided all subjects into quartile groups according to their IQ and then compared prefrontal GFC values between the 1^st^ quartile with the lower IQ scores (3 subjects, mean IQ = 113 ± 4.3) and the 4^th^ quartile with the higher IQ scores (3 subjects, mean IQ = 131 ± 1.2; the difference in IQ compared to the 1^st^ quartile is significant: t = 7.0, p < 0.01). The HbT-based connectivity maps revealed higher correlations for the majority of pairwise connections in the group with higher IQ (Fig 1). The corresponding PFC GFC values for these two groups were 0.36 ± 0.16 (the lower IQ group) and 0.69 ± 0.18 (the higher IQ group). The difference in GFC between groups was significant (t = 2.3; p < 0.05).

**Figure 1:**
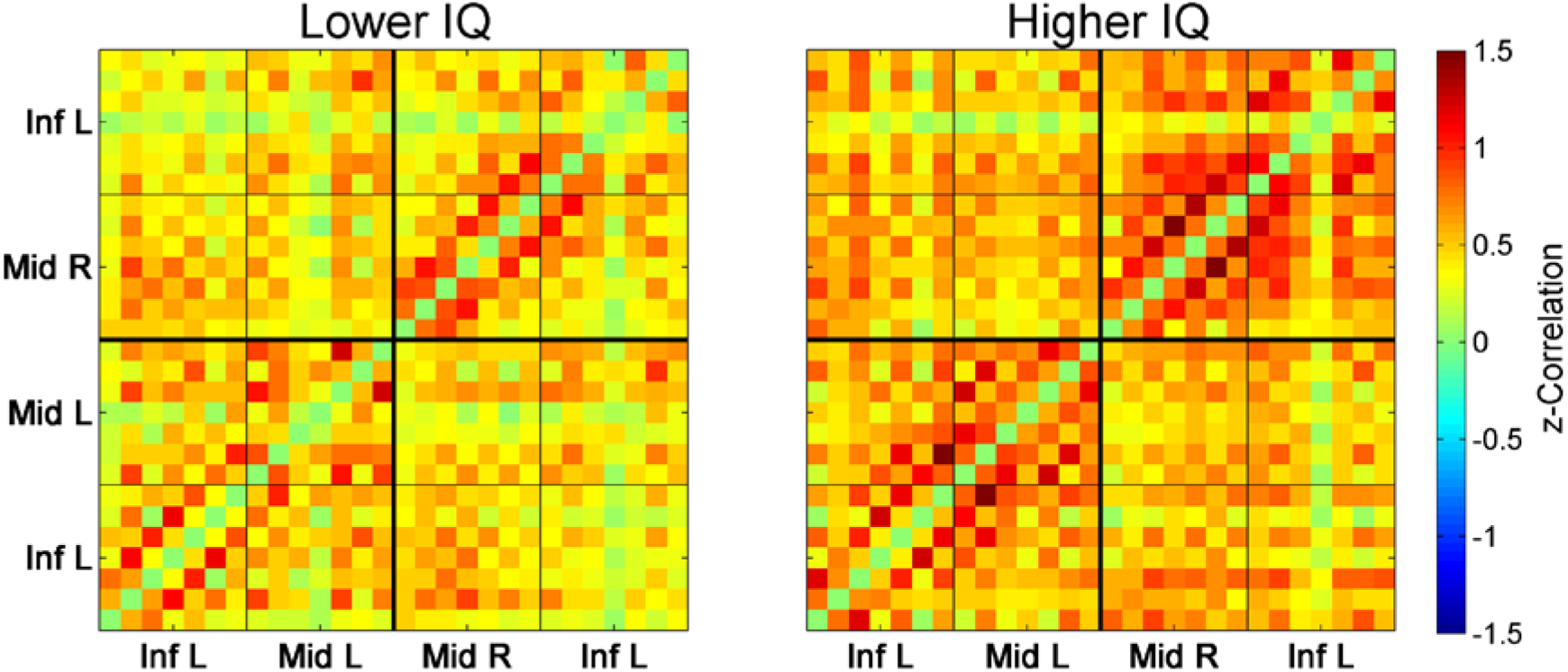
Connectivity matrices for the 1^st^ quartile (3 subjects with the lower IQ scores, *left*) and the 4^th^ quartile (3 subjects with the higher IQ scores, *right*). Each (28 x 28) matrix represents Fisher-transformed correlations calculated between all 28 optical channels pairwise. Correlation between channels v1 and v2 is represented by color at point with x-coordinate = v1 and y-coordinate = v2 (correlations for each channel to itself are intentionally zeroed and not included into the analysis). Matrices are symmetrical along the main diagonal from bottom-left to top-right. Inf L and Inf R are left/right inferior frontal gyrus (IFG), Mid L and Mid R are left/right middle frontal gyrus (MFG).

We then correlated individual GFC values with the corresponding IQ and PPT scores across all subjects. The prefrontal GFC (for all three hemodynamic signals) was found to positively correlate with the individual IQ scores with the highest significance for the total hemoglobin (Fig 2).

**Figure 2:**
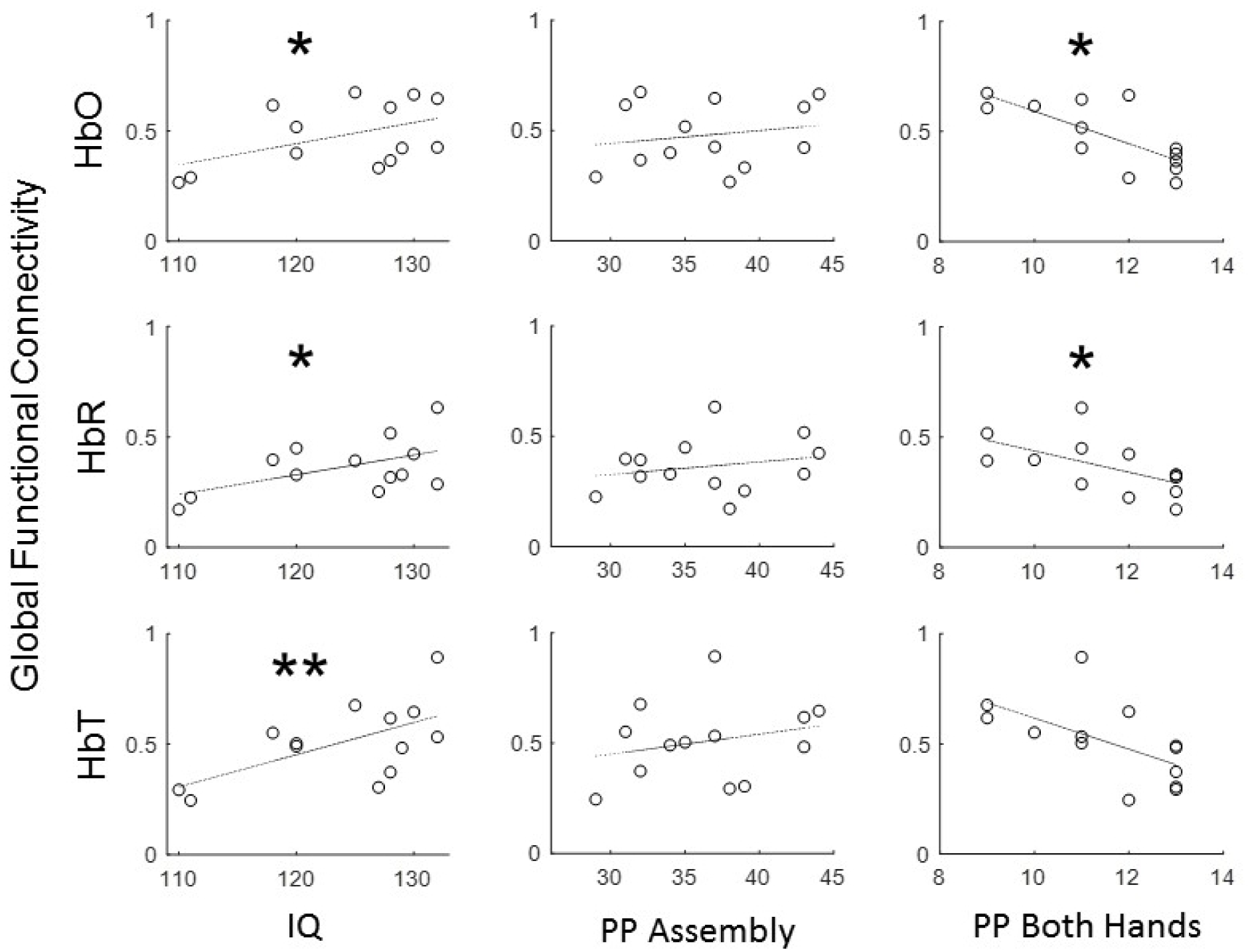
Relationship between Global Functional Connectivity (GFC) of the prefrontal cortex and the results of cognitive and behavioral tests (WASI and Purdue Pegboard Test) across a cohort of 13 healthy adult individuals. GFC showed positive correlation with IQ for all three optical signals and the lack of or negative correlation with motoric skills as measured by Purdue Pegboard Test. HbO, HbR and HbT are oxygenated, deoxygenated and total hemoglobin, respectively. PP Assembly and PP Both Hands are scores from the corresponding subtests of the Purdue Pegboard Test. Significance of linear regressions: one asterisk, p < 0.05; double asterisk, p < 0.01.

Interestingly, the prefrontal GFC did not correlate with the PP Assembly score and showed a significant negative correlation with the PP Both Hands score for HbO and HbR (Fig 2).

## 4. Discussion

The use of near-infrared spectroscopy in the studies of brain networks is growing in popularity. One limitation of NIRS is that it cannot reach deep brain structures. However, all global brain networks have large cortical components (Fox et al., 2005). This is especially true for the cognitive control network the major component of which is the frontoparietal network easily accessible by NIRS. In the previous NIRS study, we have shown that activation in PFC scales linearly with working memory load (Fishburn et al., 2014) thus confirming the adequacy of optical imaging for functional assessment of the frontoparietal network. The advantage of NIRS is that functional connectivity can be measured through temporal correlation between two raw time series with minimal preprocessing because optical signals are much less sensitive to motion (a significant problem in fMRI studies) and have higher sampling rate (50 Hz with the CW6 instrument) than fMRI (0.3-0.5 Hz). This prevents aliasing of higher frequency activity such as respiratory (∼0.2 Hz) and cardiovascular (∼1 Hz) signals into low frequency (<0.1 Hz) fluctuations through which functional connectivity is usually assessed. Higher temporal resolution of NIRS and its ability to measure relative changes in concentrations of two forms of hemoglobin separately (oxygenated, HbO and deoxygenated or reduced, HbR; their sum also determines the relative changes in tissue total hemoglobin), and this provides more information about brain hemodynamics. Along with the portability of NIRS allowing its use even in neonates at bedside and senior populations, the existing data demonstrate the capability of NIRS to provide effective tools complementary to the gold standard BOLD fMRI, to study the architecture of brain functional networks.

As Cole et al. emphasize, the most intriguing and mysterious property of the frontoparietal network is its ability to meaningfully contribute to a wide variety of task demands and moreover, being most active during the implementation of novel and non-routine tasks that the system has not had a chance to adapt to by practice or during evolution (Cole et al., 2013). These authors further demonstrated that such an extraordinary ability of the FPN results from a specific anatomical and functional organization of this system which involves *flexible hubs* i.e., brain regions that can flexibly and rapidly change their patterns of functional connectivity with other brain networks in order to achieve cognitive control across a variety of tasks (the flexible hub theory, Cole et al., 2013).

A recent study by Franzmeier et al (2017) addressed the role of FPN in the development and maintenance of cognitive reserve in Mild Cognitive Impairment (MCI), a clinical precursor of AD. Using fMRI these authors demonstrate that resting-state global functional connectivity of FPN correlates with a proxy measure of cognitive reserve (years of education) in both healthy controls and individuals with MCI and thus can be used as a potential biomarker of changes in cognitive reserve in patients at increased risk of AD (Franzmeier et al., 2017).

Our study reveals that functional connectivity of the FPN (as expressed by the GFC value) positively correlates with IQ in healthy individuals. Such correlation was observed for all forms of the optical signal (HbO, HbR and HbT) showing the highest significance of the correlation with IQ for the total hemoglobin (HbT). Although we used individual IQ scores as a measure of cognitive reserve instead of the number of years of education, our results are in good agreement with the results of Franzmeier et al (2017). Interestingly, the relationship between the PFC connectivity and the results of the Purdue Pegboard Test was quite different. Prefrontal connectivity did not correlate with the scores of the PP Assembly subtest and negatively correlated with the scores of the PP Both Hands subtest. These data can be understood and interpreted in the context of the functional role of the frontoparietal network. The FPN is the major component of the cognitive control network which is functionally distinct from other more specific networks including the sensorimotor network which is involved in motor functions. Thus, the lack of or negative correlation between PFC GFC and motoric skills (as measured by PPT) can be expected. Along with the positive correlation with IQ, these data serve as further evidence for cognitive specificity of the frontoparietal network.

In conclusion, our data provide further confirmation that cognitive reserve depends on the functional integrity of the frontoparietal network, a major cognitive control hub. Also, our experimental approach demonstrates that functional connectivity can be assessed by optical imaging using the NIRS technology. As a cost-efficient and noninvasive technology, NIRS can be used in the development of new neurophysiological measures of cognitive reserve with numerous possible applications in the context of healthy aging and cognitive disorders.

## Acknowledgment

This study was supported by awards R01EB006589 from the National Institute of Biomedical Imaging and Bioengineering and R21GM103526 from the National Institute of General Medical Sciences. The content is solely the responsibility of the author and does not necessarily represent the official views of the National Institutes of Health.

## References

Albert, S.M., Teresi, J.A., 1999. Reading ability, education, and cognitive status assessment among older adults in Harlem, New York City. Am J Public Health 89, 95–7.

Burke, D.M., Mackay, D.G., 1997. Memory, language, and ageing. Philos Trans R Soc Lond B Biol Sci 352, 1845–56.

Cole, M.W., Yarkoni, T., Repovs, G., Anticevic, A., Braver, T.S., 2012. Global connectivity of prefrontal cortex predicts cognitive control and intelligence. J Neurosci 32, 8988–99.

Cole, M.W., Laurent, P., Stocco, A., 2013. Rapid instructed task learning: a new window into the human brain’s unique capacity for flexible cognitive control. Cogn Affect Behav Neurosci 13, 1–22.

Duncan, J., 2010. The multiple-demand (MD) system of the primate brain: mental programs for intelligent behaviour. Trends Cogn Sci 14, 172–9.

Fishburn, F.A., Norr, M.E., Medvedev, A.V., Vaidya, C.J., 2014. Sensitivity of fNIRS to cognitive state and load. Front Hum Neurosci 8, Article 76 (doi: 10.3389/fnhum.2014.00076).

Fox, M.D., Snyder, A.Z., Vincent, J.L., Corbetta, M., Van Essen, D.C., Raichle, M.E., 2005. The human brain is intrinsically organized into dynamic, anticorrelated functional networks. Proc Natl Acad Sci U S A 102, 9673–8.

Fox, M.D., Raichle, M.E., 2007. Spontaneous fluctuations in brain activity observed with functional magnetic resonance imaging. Nat Rev Neurosci 8, 700–11.

Franzmeier, N., Caballero, M.A.A., Taylor, A.N.W., Simon-Vermot, L., Buerger, K., Ertl-Wagner, B., Mueller, C., Catak, C., Janowitz, D., Baykara, E., Gesierich, B., Duering, M., Ewers, M., 2017. Resting-state global functional connectivity as a biomarker of cognitive reserve in mild cognitive impairment. Brain Imaging Behav 11, 368–382.

Medvedev, A.V., Kainerstorfer, J.M., Borisov, S.V., VanMeter, J., 2011. Functional connectivity in the prefrontal cortex measured by near-infrared spectroscopy during ultra-rapid object recognition. J Biomed Opt 16, 016008.

Medvedev, A.V., 2013. Does the resting state connectivity have hemispheric asymmetry? A near-infrared spectroscopy study. NeuroImage 85, 400–7 (doi: 10.1016/j.neuroimage.2013.05.092).

Niendam, T.A., Laird, A.R., Ray, K.L., Dean, Y.M., Glahn, D.C., Carter, C.S., 2012. Meta-analytic evidence for a superordinate cognitive control network subserving diverse executive functions. Cogn Affect Behav Neurosci 12, 241–68.

Stern, Y., 2009. Cognitive reserve. Neuropsychologia 47, 2015–28.

Strangman, G., Boas, D.A., Sutton, J.P., 2002. Non-invasive neuroimaging using near-infrared light. Biol Psychiatry 52, 679–93.

Valenzuela, M.J., Sachdev, P., 2006. Brain reserve and dementia: a systematic review. Psychol Med 36, 441–54.

